# Soil microbial legacies and cultivar compatibility modulate the responses of wheat to drought

**DOI:** 10.1101/2025.09.29.679177

**Authors:** Barkha Sharma, Michel Cigan, Martin Schädler, Hamed Azarbad

**Author notes:** **Correspondence**: Department of Biology, Evolutionary Ecology of Plants, Philipps-University Marburg, Karl-von-Frisch-Strasse 8, 35043 Marburg, Germany. E-mail address.

## Abstract

Global climate change stressors are altering soil function and reducing crop yields, yet the role of soil microbial legacies in shaping plant stress responses remains poorly understood. Here, we tested how long-term farming (organic vs conventional) and climate (ambient vs future) histories of soil microbiomes influence wheat performance under drought. Soil samples were collected from long-term experimental plots of the Global Change Experimental Facility (GCEF, Germany) and used to extract microbial communities, which were then used to inoculate sterilized potting soil in which two wheat cultivars, drought-sensitive Nordkap and drought-tolerant SU Fiete, were grown under controlled greenhouse drought. Our results showed that microbial inoculation enhanced germination relative to non-inoculated, with conventional–ambient microbiomes most strongly promoting emergence, while organic–future microbiomes suppressed seed germination. Under drought, aboveground fresh biomass and dry weight content diverged by interaction between cultivar and microbial legacy in such a way that Nordkap performed best with future-climate microbiomes, whereas SU Fiete benefited from ambient-climate microbiomes. The rhizosphere of plants inoculated with organic-derived microbes harbored a larger unique ASVs, with 442 bacterial and 70 fungal ASVs, compared with 381 bacterial and 48 fungal ASVs unique to conventional-derived microbes. We further showed that rhizosphere bacterial communities were influenced by complex interactions between microbial history (farming and climate), cultivar, and water stress, while fungal communities tracked only farming history, with organic legacies buffering fungal diversity under drought. Together, these results demonstrate that soil microbiomes retain the imprint of past management and climate, and that these legacies can either buffer or exacerbate plant stress responses depending on host genotype.

## 1. Introduction

Advanced climate and agricultural models predicted a fundamental shift in global crop production by 2030 due to a projected increase in temperature, altered precipitation patterns, and elevated surface carbon dioxide concentration (Tebaldi *et al*. 2021). Over the past half century, climate-induced trends have already contributed to a decline of 4-13% in global crop yield of barley, maize, and wheat compared to projections under stable climate conditions (Lobell and Di Tommaso 2025). Increasing temperatures combined with atmospheric drying have intensified drought frequency and severity. Europe, for instance, has already witnessed severe drought events in 2003, 2010, and 2018, each resulting in major agricultural losses. In 2018 alone, Germany experienced a 17.5% reduction in winter wheat yield (Rakovec *et al*. 2022; Riedesel *et al*. 2023). Given the anticipated rise in global wheat demand (Zidi *et al*. 2025), addressing drought-induced yield losses is becoming increasingly urgent.

As sessile organisms, plants have evolved multiple strategies to tolerate, resist, or avoid the stresses imposed by water stress, including physiological adjustments, modified root system architecture, and regulation of stomatal closure in aerial tissues to minimize water loss (Tebaldi *et al*. 2021; Shoaib *et al*. 2022). These stress-induced modifications also alter the belowground rhizodepositions, which serve as the primary substrates and signaling molecules for microbial colonization, thus affecting the rhizosphere microbiome (Zhalnina *et al*. 2018; Azarbad 2024). As a highly active biological zone, the composition and functionality of root exudates are strongly shaped by biotic factors such as host genotype and its associated heritable functional traits (Li *et al*. 2023b). For instance, drought-tolerant maize cultivars maintain stable exudate profiles (e.g., fumaric acid levels) under water stress, whereas drought-sensitive cultivars exhibit pronounced shifts (Song *et al*. 2012). These genotype-specific exudation patterns chemoattract and selectively enrich microbial taxa that can either mitigate drought effects or strengthen plant stress responses (Preece and Peñuelas 2016; Li *et al*. 2023a). At the microbial community level, ecological legacies can further govern plant–microbe interactions. For instance, locally adapted microbiomes that are shaped by soil conditions and climate may determine the degree to which rhizosphere microbiomes confer host responses to stress (Azarbad *et al*. 2018, 2020; Lumibao *et al*. 2022). Repeated drought events have been shown to enrich soil and plant microbiomes for taxa with better survival traits (sporulation, exopolysaccharide production, osmolyte accumulation), creating a “stress memory” or histories that enhance tolerance to future water deficits (Azarbad *et al*. 2018; Bremer and Krämer 2019; Schmidt *et al*. 2024). These stress-experienced microbes carry certain functions, such as genes for carbohydrate and amino acid metabolism and transport, which are critical for surviving episodic drought stress (Xie *et al*. 2021) that can act as inocula to improve plant performance under subsequent droughts (Lau and Lennon 2012; Vilonen *et al*. 2023).

Agricultural management practices have been known to shape soil microbial communities. As an example, organic farming, which is characterized by organic fertilization, crop rotations, and the absence of synthetic pesticides (Azarbad 2022), often supports higher soil microbial diversity and more heterogeneous microbial communities (Lupatini *et al*. 2017; Nam, Lee and Choi 2023; Lori *et al*. 2024). In contrast, intensified agricultural practices such as conventional farming, with mineral fertilizers and pesticide use, simplify microbial networks, reduce the abundance of keystone taxa, and lower soil functionality (Banerjee *et al*. 2019; van Rijssel *et al*. 2025). Global climate change is likely to amplify the negative impacts of agricultural intensification on various ecosystem functions (IPBES, 2019). However, we have a limited understanding of how agricultural fields managed conventionally or organically with contrasting exposure to ambient and future climate scenarios (drought and warming) withstand drought stress. Understanding how different plant genotypes select and interact with soil microbiomes with different farming and climate histories can help to adjust agricultural management to best support or maintain an adaptive soil and plant-associated microbiome to increase crop production under drought.

Here, we investigated whether soil microbiomes shaped by contrasting agricultural and climatic histories can enhance wheat performance under drought. For that, we collected soils from the Global Change Experimental Facility (GCEF), a long-term field site in Germany that imposes ambient versus future-climate (warming and altered precipitation patterns) treatments across conventional and organic managed plots. We first extracted microbial communities from each soil history and used these inocula to inoculate sterilized potting soil. Two wheat cultivars (SU Fiete, a drought-tolerant cultivar; Nordkap, a drought-sensitive cultivar) were then grown in the inoculated and non-inoculated control soil under a controlled greenhouse drought experiment. We monitored germination and aboveground biomass, and studied rhizosphere microbiomes using 16S and ITS amplicon sequencing. With this experimental setup, we aimed to answer the following questions: do the agricultural or climatic legacies of soil microbiomes influence the response of two different wheat cultivars under water stress, and how do soil microbiome legacies restructure rhizosphere communities? We hypothesized that organic farming legacies would support higher microbial diversity, whereas future-climate legacies would enrich for stress-adapted microbes. We further predicted that cultivar-specific responses determine whether microbial legacy would buffer or exacerbate wheat responses to drought. Specifically, we expected that the drought-sensitive cultivar would become selective in the rhizosphere, thus recruiting microbial partners capable of compensating for its physiological limitations, whereas the drought-tolerant cultivar would maintain broader compatibility with diverse microbes.

## 2. Materials and methods

### 2.1. Soil sampling

The soil samples were provided by GCEF, a large multi-year field experiment in Saxony-Anhalt, Germany (51° 23’ 30N, 11° 52’ 49E, 116 m a.s.l.), which is designed to simulate the changing global climate conditions to examine the effect of different ecosystem processes across various land-use types and intensities. The field consisted of 50 plots (16 m x 24 m each) arranged into 10 main plots, with 5 sub-plots per main plot (Schädler *et al*. 2019). The different land-use regime applied to these 5 plots are (a) conventional farming: mineral fertilizers and pesticides applied as per conventional agricultural practices, (b) organic farming: mechanical weed control, organic fertilization, untreated seeds, and pesticide are not used, (c) intensively used grassland: frequent mowing to maintain high-intensity grassland usage, (d) extensively used grassland (mowing): moderate mowing to maintain lower-intensity grassland, and (e) extensively used grassland (grazing): moderate sheep grazing with same plant species composition as regime (d). In conventional farming, the crop rotation consists of winter rape, winter wheat, and winter barley, with the use of synthetic fertilizers and pesticides (see Schädler et al. (2019) for detailed information). Additionally, nitrogen is applied several times using calcium ammonium nitrate at a total rate of 40–60 kg/ha. Potassium is added as potassium chloride at 110 kg/ha, and phosphorus is supplied annually as superphosphate at a rate of 30 kg/ha. In organic systems, winter rape is rotated with legumes such as alfalfa and white clover to support nitrogen fixation naturally. Additionally, organic fields are treated yearly with 120 kg/ha of potassium from patent kali and 45 kg/ha of phosphorus from rock phosphate.

To simulate future climate conditions, half of the experimental blocks (5 blocks) were placed under a steel structure equipped with a mobile roof, side panels, and an irrigation system. By closing the roof and side panels, night temperatures were elevated, and summer precipitation was reduced by approximately 20%, while spring and autumn precipitation were increased by 10% via irrigation. This setup aligns with climate change projections for central Germany. The other 5 blocks, used as controls, were housed under a similar structure without interventions to maintain ambient climate conditions. In this study, we collected soil from the GCEF field from the following treatment combinations of farming type and climate condition: (1) conventional farming under ambient climate (CA), (2) conventional farming under future climate (CF), (3) organic farming under ambient climate (OA), and (4) organic farming under future climate (OF). These are hereafter referred to as soil history types. Twenty soil sub-samples (2 farming histories × 2 climate histories × 5 replicate plots) were collected from the topsoil (0–15 cm, ∼0.5–0.6 kg per replicate), thoroughly homogenized within each treatment, and pooled to generate one representative composite sample per soil history type (CA, CF, OA, OF). All samples were stored at 4 °C prior to microbial extraction.

### 2.2. Extraction of soil microbiome

Microbial extractions were conducted for each soil history type (CA, CF, OA, OF) using sterilized equipment. For each extraction, 15 g of soil was mixed with 150 ml of Milli-Q water in an Erlenmeyer flask and shaken for 1.5 hours at 300 rpm to release the microbial community, leaving behind small stones and woody debris. The liquid portion was then transferred to 50 ml centrifuge tubes and centrifuged at 22 °C, 1500 rpm for 5 minutes. The supernatant was collected in glass bottles and stored at 4 °C overnight. Our previous study demonstrated that the microbial extraction protocol preserves community composition in such a way that microbial profiles in the suspensions match closely those in the source soils prior to extraction (Ornik *et al*. 2024). 300 g of sterilized potting soil (HAWITA Fruhstorfer®; HAWITA Gruppe GmbH) was placed into each of 100 pots (11 cm x 11 cm x 21.5 cm, tapering to 8.5 cm x 8.5 cm at the base). Soil sterilization was achieved by autoclaving twice: first at 121 °C for 60 minutes, then again under the same conditions 24 hours later. Each soil microbial extract (80 ml) was used to inoculate 40 pots, while 40 control pots received 80 ml of Milli-Q water. To ensure even microbial distribution, soils were mixed with sterile tubes. Pots were then placed in the greenhouse for two weeks to allow microbial growth and acclimation before sowing seeds. It is important to note that on the same set of samples, we have previously shown that soil from conventional plots contains higher levels of nitrogen and phosphorus (Ornik *et al*. 2024). However, in this experiment, soil extracts represented only a tiny fraction of the total soil volume (80 ml suspension into 300 g sterile potting soil), making any nutrient contribution both highly diluted and transient.

### 2.3. Experimental design

The pot experiment was conducted at the Phillips Universität Marburg Greenhouse (50.8083° N, 8.7694° E), with controlled light conditions (Fig. 1A). After two weeks of microbial incubation, wheat seeds of the cultivars Nordkap and SU Fiete were planted. For each soil type (with farming and climate history) and uninoculated controls, half of the pots were sown with Nordkap and the remaining half with the SU Fiete cultivar. Six seeds were planted 5 cm deep per pot. The setup was placed under Neusius LED grow lights (Neusius Pflanzenlicht, Item-Nr. LED/D-E4-180VR) (12-hour photoperiod, day-night temperature) in a greenhouse. The pots were watered with 100 ml of tap water every two days for two weeks to ensure consistent initial growth conditions. Germination rate and time were measured over a period of 6 days. Germination rate was calculated as the percentage of seeds that successfully sprouted each day relative to the total number of seeds per pot (6 seeds). After 14 days, when seedlings were established, plants were thinned to five plants per pot and watering conditions were adjusted, where water stress (drought) was then applied to half of the pots in each soil and cultivar group, with drought-treated pots receiving 30 ml of water compared to 100 ml for well-watered control plants. The watering schedule was adjusted based on temperature and upper soil moisture to maintain drought conditions for six more weeks. Pots were arranged in a fully randomized design, and their positions were re-randomized weekly. The experimental setup resulted in the total combination of 20 treatments as follows: 4 microbial extracts (2 farming × 2 climate histories) plus a not-inoculated control × 2 wheat cultivars × 2 water stress treatments (drought plus well water control). Each treatment was replicated 5 times in a randomized design (20 treatments × 5 replicates = 100 pots). Plants were sampled for the rhizosphere sampling and measurement of both fresh and dry weight aboveground biomass. Dry weight was determined by drying the biomass at 60 °C for 4 days, until a constant weight was achieved. Dry weight content (DWC) was calculated by dividing the dry biomass by the fresh biomass and multiplying the result by 100, and expressed as a percentage.

**Fig 1.**
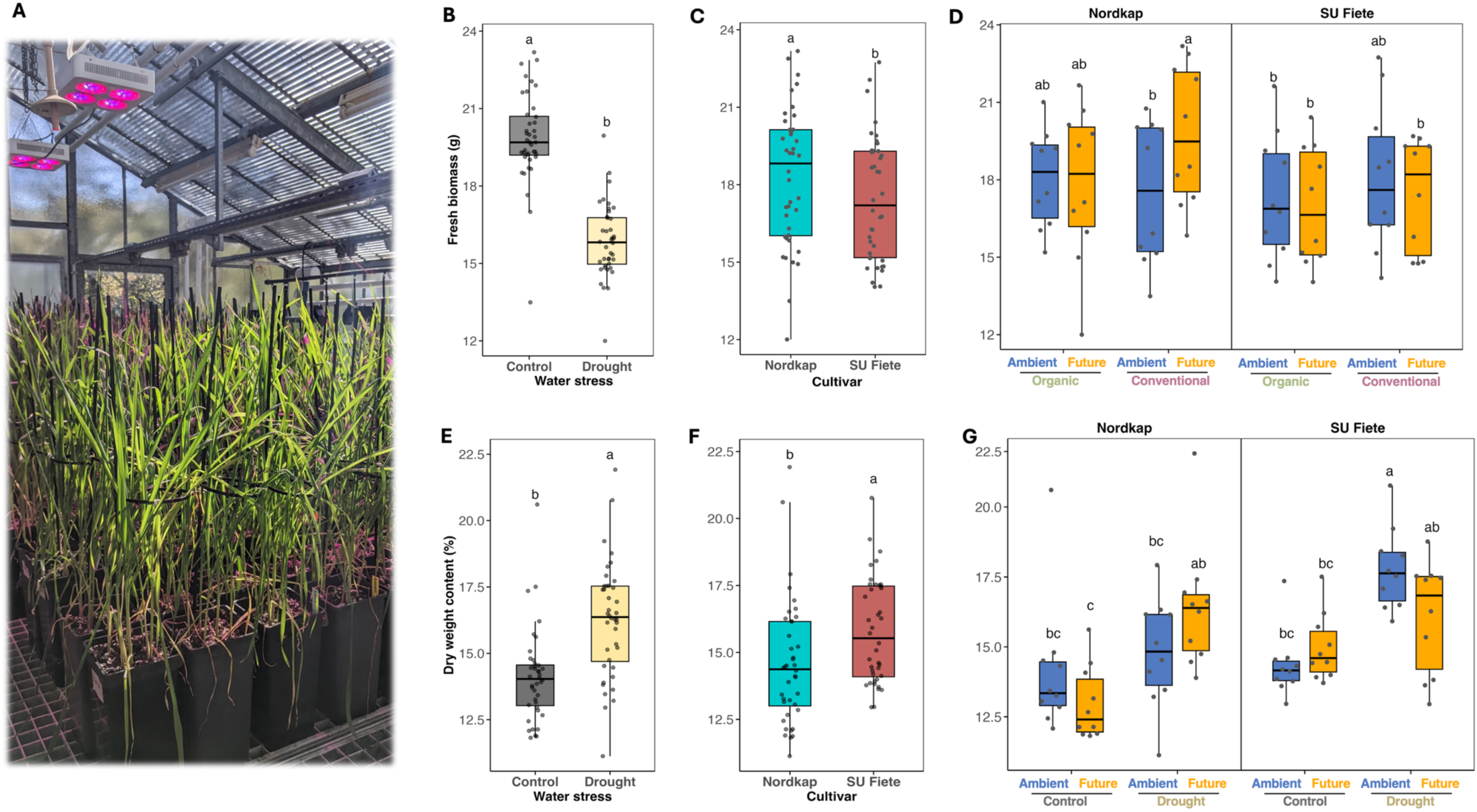
(A) Greenhouse setup of the wheat drought experiment. Pots were filled with sterilized potting soil and inoculated with microbial extracts from soils differing in farming (organic vs conventional) and climate (ambient vs future) histories at the Global Change Experimental Facility (GCEF, Germany). Two wheat cultivars (Nordkap and SU Fiete) were grown under controlled conditions and subjected to either well-watered or drought treatments for eight weeks. Fresh biomass variation by (B) water stress, (C) cultivar type, and (D) significant interaction terms. Dry weight content variation by (E) water stress, (F) cultivar type, and (G) significant interaction terms. Letters represent significant differences based on *post hoc* Tukey’s test.

### 2.4. DNA extraction, amplicon sequencing and data processing

For the extraction of DNA, rhizosphere soil was collected from all plants in each pot within each treatment. We used DNeasy® PowerSoil® Pro Kit (QIAGEN) to extract DNA from soil based on the manufacturer’s protocols. DNA samples were sent to Novogene (UK) for libraries preparation and Illumina MiSeq (paired-end) sequencing. For the bacterial 16S rRNA gene, the V5–V7 region was amplified using primers 799F (AACMGGATTAGATACCCKG) and 1193R (ACGTCATCCCCACCTTCC). As for fungi, ITS1 was amplified based on ITS1-1F-F (CTTGGTCATTTAGAGGAAGTAA) and ITS1-1F-R (GCTGCGTTCTTCATCGATGC). Processing sequence data was conducted in QIIME 2 to generate Amplicon Sequence Variants (ASVs) for both bacterial (16S rRNA) and fungal (ITS) datasets. Briefly, primer sequences were removed, and reads were truncated ensuring that only high-quality bases remained prior to denoising. Chimeric sequences were identified and removed with the removeBimeraDenovo function of DADA2. For taxonomic affiliations of the resulting amplicon sequence variants (ASV), a naive Bayesian classifier was performed based on the SILVA database v138. ASVs unassigned at the phylum level, together with chloroplast and mitochondrial reads, were removed from the dataset. Before further analysis, the following filtering criteria were applied: samples should have more than 100 ASV reads, and any ASVs with less than five reads in a given sample were removed. Furthermore, any ASV that was found in only one sample was discarded from the data set. For the calculation of alpha diversity, the data were normalized on the basis of sequencing depth. For beta diversity, we normalized the data set based on the relative abundance of ASVs in each sample (Walsh *et al*. 2021). After filtering out ASVs occurring in less than 2 samples and less than 100 reads, we identified a total of 9343 bacterial ASVs and 1626 fungal ASVs in the rhizosphere.

### 2.5. Statistical analysis

Statistical analyses were performed using R (Version 4.3.2; R Development Core Team, 2024) and R Studio (Version 2023.12.1+402). We first tested the influence of soil history-derived microbial communities on plant growth parameters, including germination rate, fresh biomass, and DWC. For that, ANOVAs were conducted to evaluate variations in plant performance across experimental factors, including farming practices, climate type, cultivar, and water stress treatment. Tukey’s (HSD) test was used for post-hoc comparisons among experimental factors. To isolate the effect of microbial inoculation, we calculated the difference (Δ) in fresh biomass and DWC between inoculated and non-inoculated plants for each combination of farming history, climate history, and watering treatment. The *α-*diversities of the rhizosphere soil microbiome were calculated using ASV richness and Shannon-index (Shannnon and Weaver 1949) via *agricolae* package (Mendiburu and Simon 2009). ANOVAs test was performed to evaluate the direct and interactive effects of experimental factors on diversity indexes. Next, we conducted Principal Coordinate Analyses (PCoA) using the vegan package (Oksanen *et al*. 2022) to visualize microbial community composition and the Permanova test (through the ‘*adonis2*’ function) to evaluate the effect of experimental factors on the microbial community structure. Our analysis was based on the relative abundance of ASVs, using Bray–Curtis dissimilarity. Additionally, we compared the number of shared and unique ASVs between the rhizosphere of plants grown in soil inoculated with soil microbial extracts and non-inoculated control plants and plotted results as *UpSetPlot* using *ComplexUpset* package (Lex *et al*. 2014; Krassowski 2020). Further, we conducted ANOVA to analyze the impact of each experimental factor on the relative abundance of dominant microbial taxa (mean relative abundance of >1%) at the genus taxonomic level. Stacked bar plots were generated to visualize the variation in the relative abundance of the dominant bacterial and fungal classes that showed significant changes due to experimental factors (based on ANOVA results). Finally, Spearman’s rank correlation (p < 0.05, Benjamini-Hochberg corrected) was applied to investigate the correlation between DWC and bacterial genus abundances (top 200 most abundant) and fungal genus abundances (all genera) under drought conditions for Nordkap and SU Fiete. Significant correlations (p < 0.05) were selected for further visualization. All plots were generated using *ggplot2* (Wickham 2016), unless specified otherwise. All statistical tests were conducted with a significance threshold of p < 0.05, and multiple testing corrections were applied where necessary.

## 3. Results

### 3.1. Plant growth parameters

The farming history of the microbial extract was the primary driver of variation in wheat seed germination (F = 54.85, Day 5 in Supplementary Table S1). Seeds inoculated with microbes originating from conventional farming exhibited the highest germination rates (day 5, Fig. S1B). Furthermore, the microbiome extract from soil exposed to an ambient climate under a conventional farming history (for both cultivars) not only promoted the fastest germination (Day 4) but also fostered the highest germination rate by Day 5 compared to other treatments (Fig. S1A-B). In contrast, organic - future inocula resulted in the lowest and slowest germination rates across all treatment combinations (Fig. S1A-B). Among the two wheat cultivars, Nordkap displayed a stronger response to microbial inoculation, especially under ambient climate and conventional farming, achieving a significantly higher germination rate (100% germination rate by Day 5) than SU Fiete (Fig. S1B, Table S1).

The ANOVA results of plant biomass revealed significant main effects of water stress (Drought vs. Control) and cultivar (Nordkap vs. SU Fiete) on fresh biomass and DWC (Table 1). As shown in Fig. 1B, across treatment combinations, water stress exposure reduced fresh shoot mass by 25.16% relative to well-watered controls. Moreover, the two cultivars showed differential responses towards fresh biomass production, with Nordkap showing a 4.11% increase compared to SU Fiete (Fig. 1C). Fresh biomass was also shaped by a three-way interaction of Farming × Climate × Cultivar (P < 0.05, Table 1). For Nordkap, plants grown in soil inoculated with microbes from future climate conditions showed higher biomass (12.93%) compared to the ambient, particularly under conventional farming, while SU Fiete responded similarly to the climate–farming treatment (Fig. 1D). Water stress led to a higher DWC, indicating reduced tissue moisture (Fig. 1E), but the magnitude of the increase depended strongly on plant cultivar and microbial climate history (Climate × Cultivar × Water stress interaction effect; Table 1; Fig. 1F and G). For instance, under water stress, SU Fiete plants grown in the soil inoculated with ambient-climate microbes showed the highest DWC values (17.78 %) compared to other treatments. However, in the case of Nordkap, plants grown in soil with future-climate inocula tended to have higher DWC values under water stress than plants exposed to ambient-climate inocula (Fig. 1G).

### 3.2. Inoculation-induced shifts in fresh biomass and dry weight content under drought

Next, to have a better picture of microbial inoculation effect on the change in fresh biomass and DWC, we compared inoculated plants versus corresponding non-inoculated plants for each treatment separately (Fig. S2; Table S2). The fresh biomass of Nordkap inoculated with microbes from future climate conditions exhibited higher fresh biomass under well water conditions, exceeding that of the control group, irrespective of the farming histories of microbes (Fig. S2A). In contrast, SU Fiete inoculated with future-climate microbial extracts under well-watered control conditions showed a significant reduction in fresh biomass compared to its non-inoculated counterpart (Fig. S2A). In the case of DWC, inoculation generally increased DWC relative to non-inoculated controls, but the direction and magnitude of responses were cultivar- and climate-dependent (Fig. S2B). In Nordkap, all microbial inocula elevated DWC, with the strongest effect under drought in combination with future-climate inocula (median Δ ≈ +3%). In contrast, SU Fiete grown in soil with future-climate inocula showed a reduction in DWC under drought relative to other treatments (Fig. S2B).

### 3.2. Microbial diversity and community structure of wheat rhizosphere

Water stress exerted a significant direct effect on bacterial richness (F = 14.11, P < 0.001), influencing both wheat cultivars (Table 2, Fig. 2A). In Nordkap, richness remained stable between well-watered and drought treatments when inoculated with future-climate microbiomes, but showed a decline under drought when paired with ambient-climate inocula, respective to the well-watered control (Fig. 2A; Table 2). On the other hand, SU Fiete exhibited greater variability across treatments, with bacterial richness being particularly reduced under drought (Table 2, Fig. 2A) and lowest under drought in the rhizosphere of plants grown in the soil inoculated with microbes from future climate (interactive effect of Climate × Cultivar × Water Stress, Table 2, Fig. 2A). Fungal richness was significantly higher in rhizosphere samples of plants inoculated with microbes of organic farming (P = 0.001; Table 2, Fig. 2C). Furthermore, the results of Shannon diversity indicated that rhizosphere of Nordkap plants grown in soil inoculated with organic farming microbes had significantly higher bacterial diversity (5.2 ± 0.34) than that of plants inoculated with inocula from conventional field (4.7 ± 0.51). However, no significant difference was observed for the SU-Fiete cultivar (significant interactive effect of Farming × Cultivar; Table 2; Fig. 2B). ANOVA revealed a significant Farming × Water Stress interaction (Table 2), indicating that drought’s effect on fungal Shannon-diversity depended on the agricultural history of the inoculated microbiome. Under well-watered (control) conditions, organic-derived inocula supported a higher Shannon index (1.91 ± 0.39) than conventional-derived inocula (1.65 ± 0.28; Fig. 2C). When subjected to drought stress, however, organic soils maintained their fungal α-diversity (1.94 ± 0.27), while diversity in conventional soils declined to 1.89 ± 0.35 (Fig. 2D).

**Fig 2.**
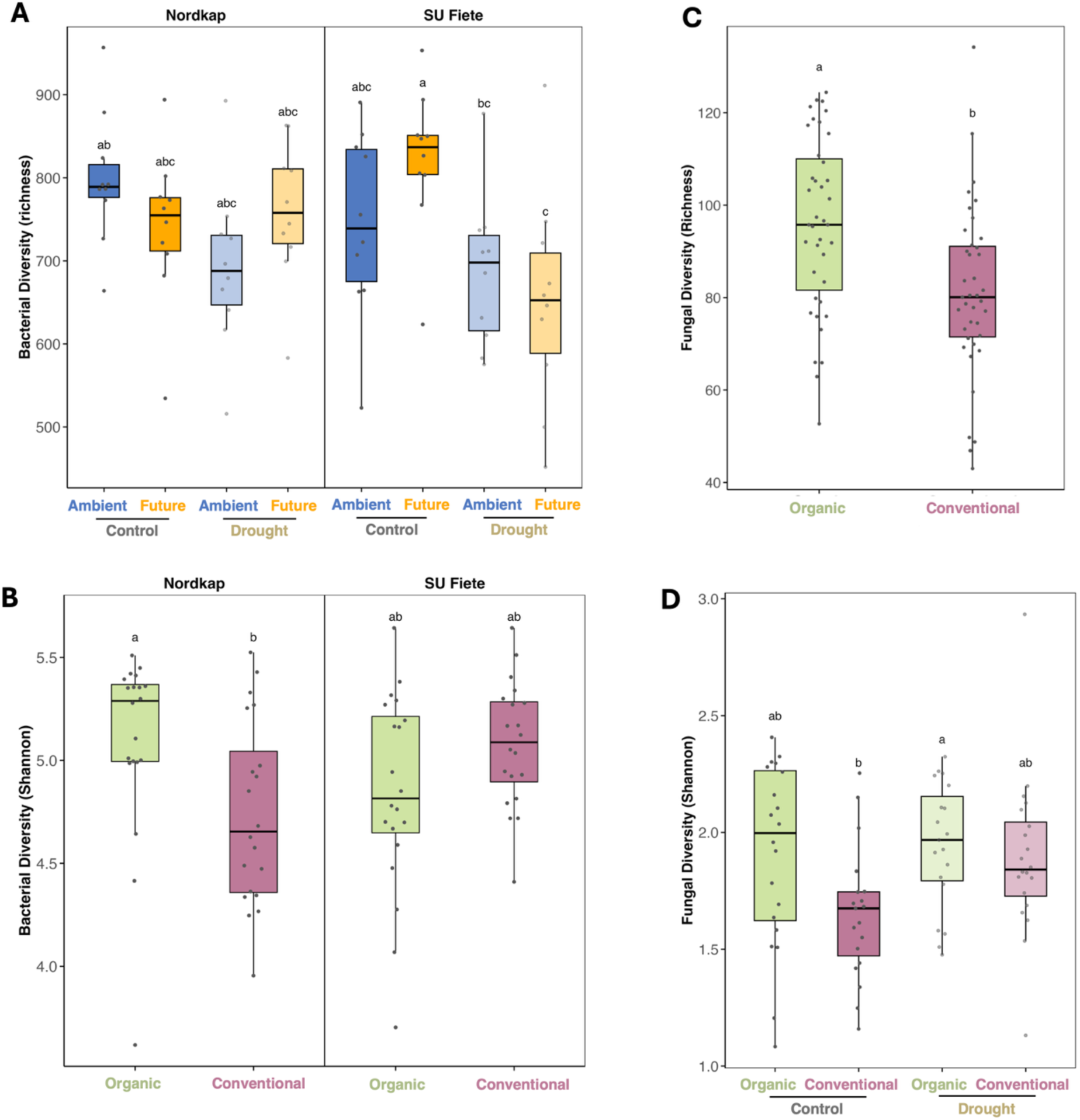
Alpha-diversity of bacterial and fungal communities in the wheat rhizosphere. (A) Bacterial ASV richness, (B) bacterial Shannon diversity (C) fungal ASV richness and (D) fungal Shannon diversity. Different letters denote statistically significant differences based on post hoc Tukey’s test.

To identify the possible effect of experimental factors on bacterial and fungal community structures, we further performed a principal coordinate analysis (PCoA) based on the Bray-Curtis dissimilarity (Fig. 3). The PCoA of rhizosphere bacterial communities showed significant (P < 0.001) effects on farming, climate, and water stress on bacterial composition, each explaining 7.27%, 3.12%, and 2.25% of variation, respectively (Table 2; Figs. 3A-C). The bacterial communities were found to be clustered more after the application of microbes with distinct farming (Fig. 3A) and climate histories (Fig. 3B). Interestingly, the main effects were further modified by a significant four-way interaction involving Farming, Climate, Water stress, and Cultivar (P < 0.01) to shape the bacterial community in the rhizosphere. Permanova analysis of rhizosphere fungal communities revealed that farming practice (organic vs conventional) was the only driver of variation of community structure (Table 2; Fig. 3D).

**Fig 3.**
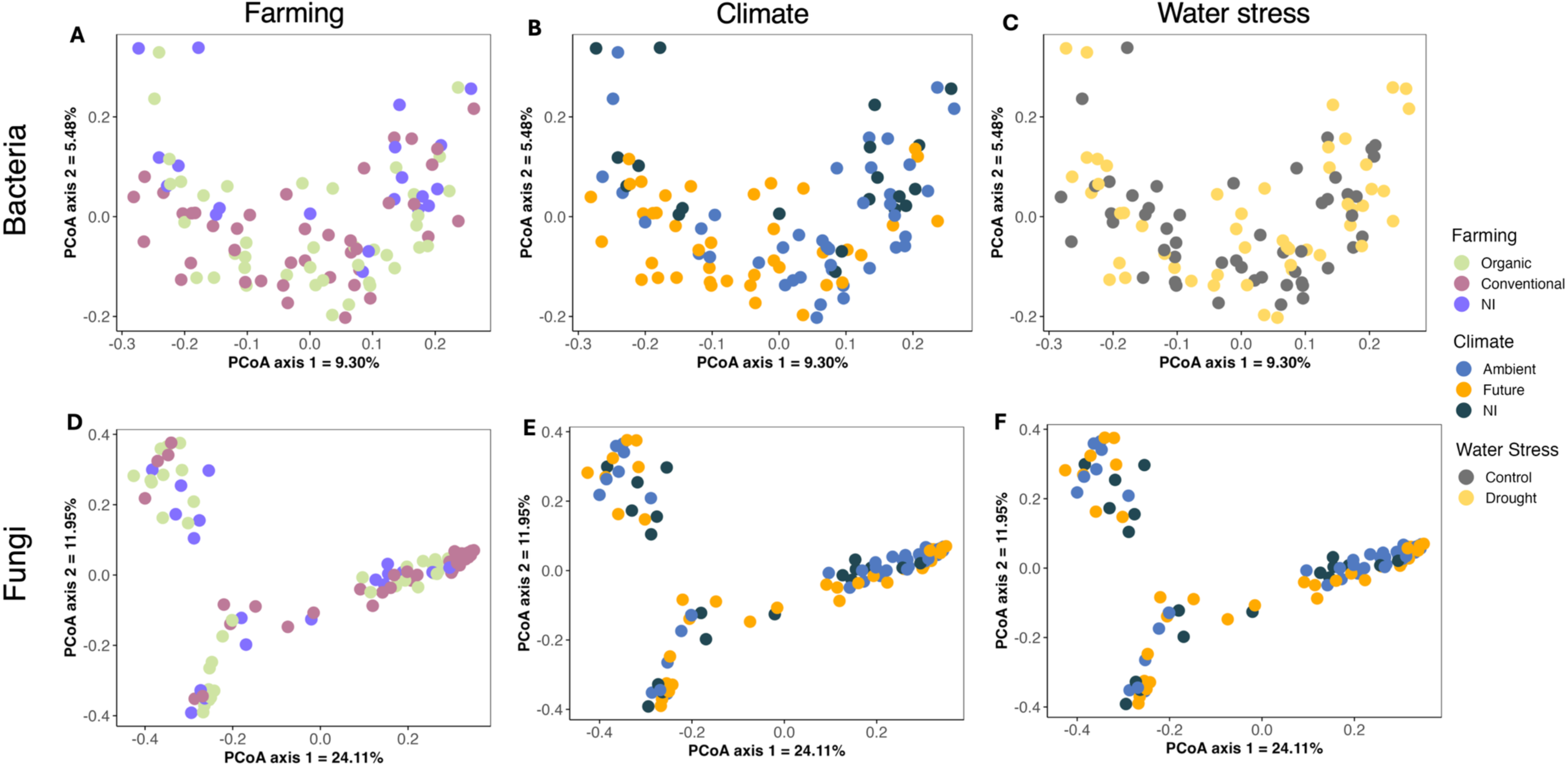
Community composition analysis of the rhizosphere microbiome of wheat. A-F, Principal coordinate analysis (PCoA) ordination plot based on Bray-Curtis dissimilarities of bacteria microbiome colored by farming system (A), Climate conditions (B), and water stress (C), and fungal microbiomes colored by farming system (D), climate conditions (E), and water stress (F), as indicated in the legend. Each data point represents a single replicate of the treatment.

### 3.3. Unique and shared ASVs between the rhizosphere of wheat plants inoculated with soil microbial extract with different farming legacies

Given that farming history emerged as the main driver of both bacterial and fungal community structure in our Permanova (Table 2) and PCoA analyses (Fig. 3), we quantified the unique and shared bacterial and fungal ASVs of the rhizosphere of plants inoculated with organic and conventional soil inocula, as well as non-inoculated control plants. Across all treatments, a core microbiome of 1,734 bacterial (Fig. 4A) and 321 fungal (Fig. 4B) ASVs colonized wheat rhizosphere irrespective of farming origin, reflecting a shared baseline of the rhizosphere-associated taxa. Beyond this core, the rhizosphere of plants grown in soil inoculated with soil microbial extract from an organic management field contributed the largest pool of unique ASVs, including 442 unique bacterial and 70 unique fungal ASVs, surpassing those samples related to conventional farming (381 bacteria; 48 fungi) and non-inoculated plants (73 bacteria; 60 fungi).

**Fig 4.**
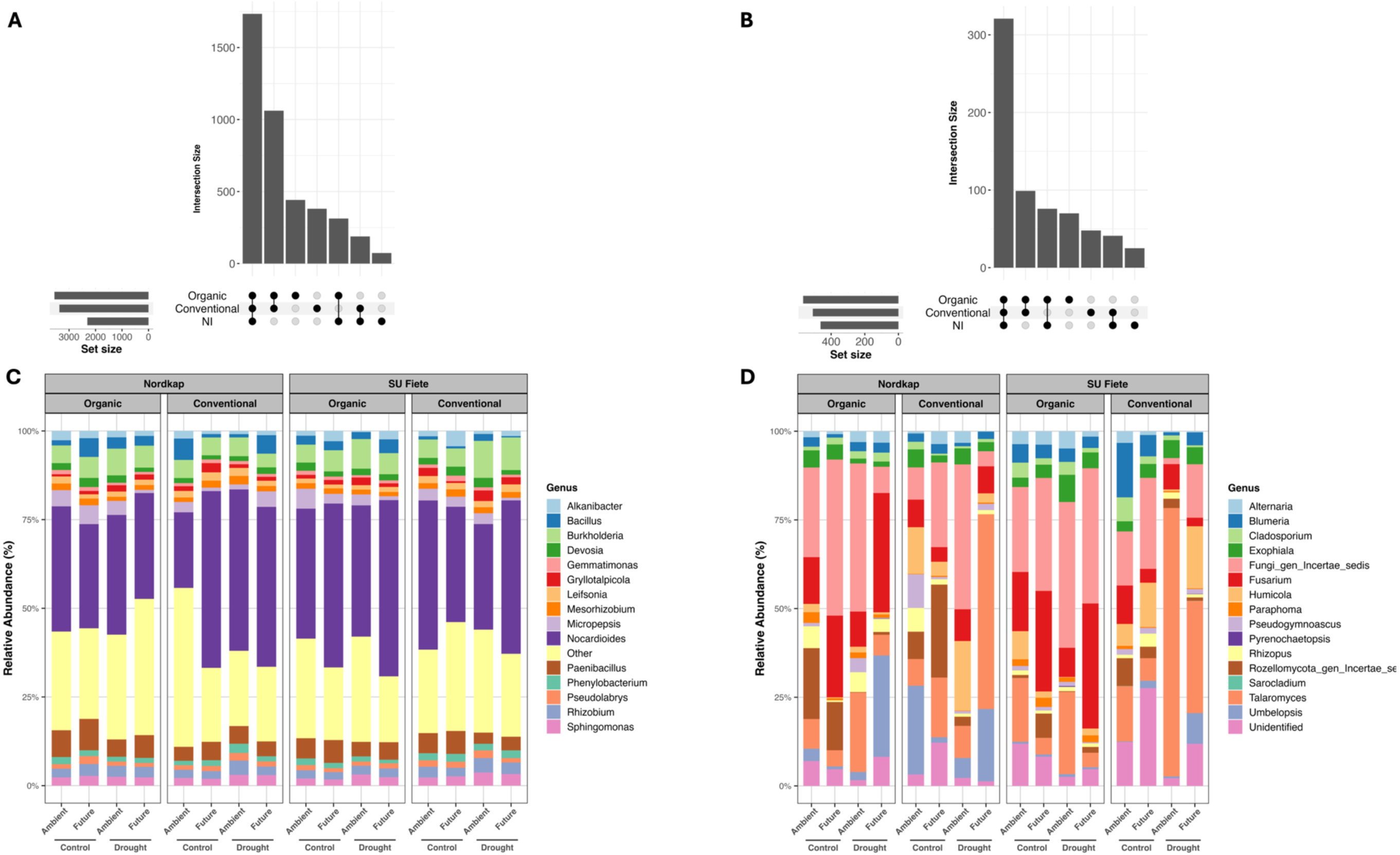
Upset plots showing unique and shared ASVs for (A) bacteria and (B) fungi between the farming system (organic vs. conventional) and non-inoculated control plants. Set size bar representing the total ASVs per group, dark circle indicates the unique ASVs, dark circle connecting bar indicates the sharing between the treatments. Relative abundance of the top 15 most dominant (C) bacterial and (D) fungal genera in the wheat rhizosphere under water stress. The top 15 most abundant genera with significant interaction are shown.

### 3.4. Community composition of the bacteria and fungi

We examined changes in the composition of bacterial (Fig. 4C; Table S3) and fungal (Fig. 4D; Table S4) communities at genus levels in rhizosphere samples from Nordkap and SU Fiete plants under experimental conditions. Across treatments, the rhizosphere was dominated by *Nocardioides*, which constituted the largest single genus in every panel of Fig. 4C and peaked in Nordkap plants inoculated with conventional–future microbiomes (Farming × Climate × Cultivar; Table S3). Climate history significantly shifted the relative abundance of *Rhizobium,* with ambient-climate inocula supported higher relative abundance than future-climate inocula (Fig. 4C; Table S3). Independent of legacy and cultivar, water stress reduced *Paenibacillus* (P<0.001) and *Sphingomonas* (P<0.001) (Fig. 4C; Table S3). Under drought, *Burkholderia* was more abundant in the rhizosphere of SU Fiete when inoculated with conventional versus organic microbiomes (Farming × Water stress × Cultivar, Table S3), an effect not observed in Nordkap. We also found that, unlike other genera, *Alkanibacter, Bacillus, Devosia,* and *Micropepsis, Phenylobacterium* and *Pseudolabrys* were significantly influenced by the interaction of farming, climate, cultivar, and water stress (four-way interaction effect; Table S3).

As for fungi, farming significantly shifted *Cladosporium*, *Fusarium*, and *Humicola* in such a way that *Humicola* was significantly higher under conventional legacies, whereas *Cladosporium* and *Fusarium* were higher under organic legacies (Fig. 4D; Table S4). Climate showed a direct effect only for *Fusarium* (P < 0.05), with higher relative abundance under organic farming (significant Farming × Climate interaction), and future-climate legacies exerted higher *Fusarium* than ambient (Fig. 4D; Table S4). The relative abundance of *Talaromyces* was significantly higher under drought stress, while *Blumeria*, *Rhizopus* (belong to *Mucoromycota* phyla), and *Rozellomycota gen Incertae sedis* were more abundant in well-watered plants. In addition, *Humicola* and *Sarocladium* reached the highest abundances under conventional–ambient microbiomes in well-watered controls, particularly in SU Fiete (significant Farming × Climate × Water stress; Table S4; Fig. 4D).

### 3.5. Correlation between microbial responders and plant growth parameters

To assess links between plant performance and rhizosphere microbiomes, we performed Spearman’s rank correlations between DWC and the 200 most abundant bacterial genera as well as all fungal genera. Because the most pronounced DWC differences occurred in Nordkap and SU Fiete under ambient-versus future-climate inocula during drought (Fig. 1G), analyses focus on these treatments. In drought-stressed Nordkap, higher DWC with future-climate inocula (Fig. 1G) coincided with a greater number of positive correlations, including associations with *Steroidobacter*, *Chthoniobacter*, and *Achromobacter* (Fig. 5A). Under ambient-climate inocula, correlations were fewer and weaker. In contrast, SU Fiete exhibited higher DWC with ambient-climate inocula, accompanied by stronger positive correlations with genera such as *Mucilaginibacter*, *Pedosphaera*, and *Puia* (Fig. 5B). Future-climate inocula also yielded positive associations in SU Fiete, but these were fewer and of lower magnitude compared with ambient-climate inocula. Across both cultivars, fungal correlations with DWC were less frequent and weaker than bacterial correlations (Table S5).

**Fig 5.**
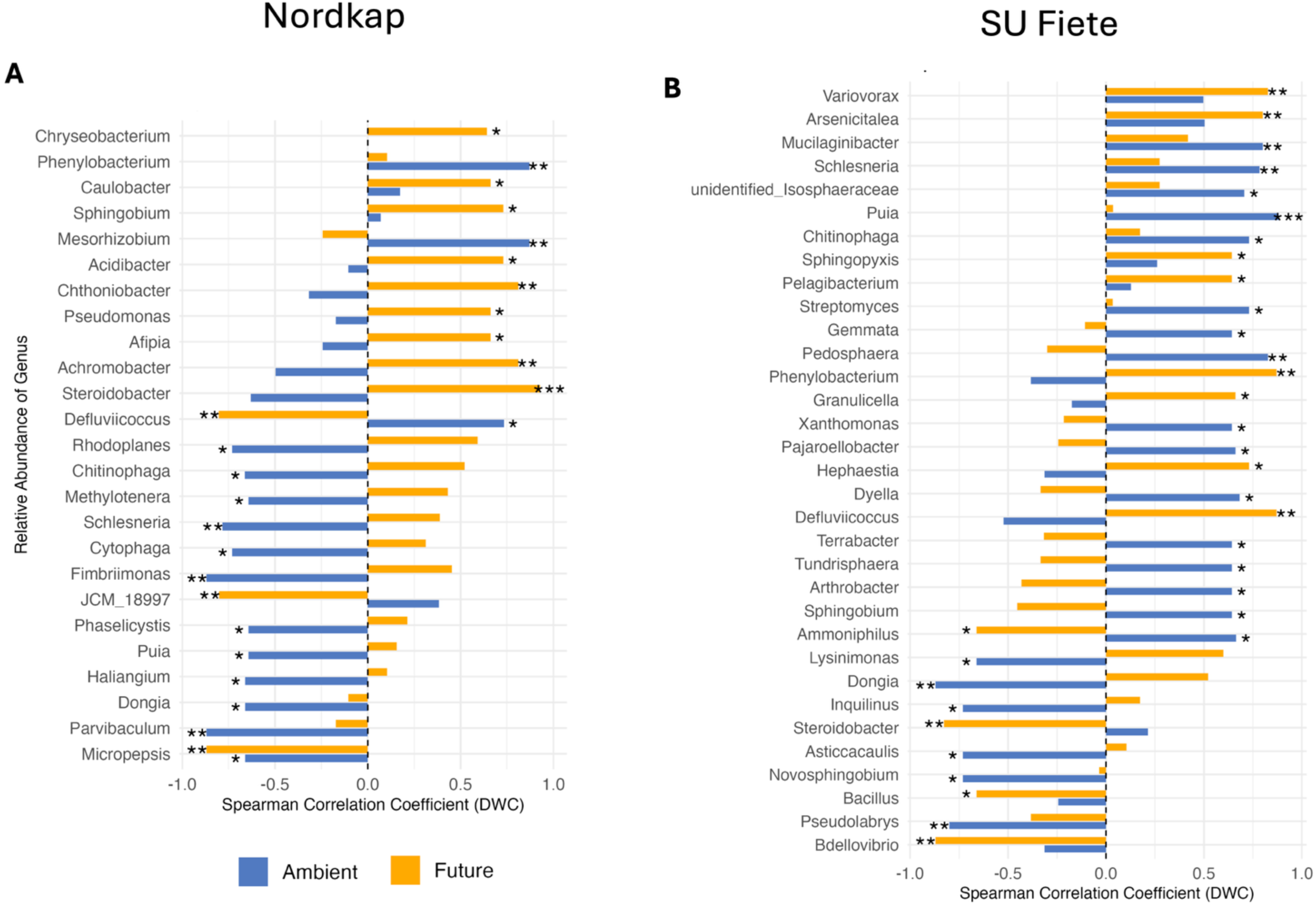
Spearman correlation between dry weight content (DWC) and bacterial abundancies in the rhizosphere of wheat under ambient and future climate conditions. (A) Correlation barplot of significant correlations (p<0.05) between DWC and the bacterial abundancies at the genus level of drought-affected Nordkap rhizosphere under ambient and future climate. (B) Correlation barplot of significant correlations (p<0.05) between DWC and the bacterial abundancies at the genus level of drought-affected of the SU Fiete rhizosphere under ambient and future climate. The color of bars indicates climate conditions: ambient (blue), future (orange). Asterisks represent significance levels (*: p < 0.05, **: p < 0.01, ***: p < 0.001 respectively).

## 4. Discussion

Our study demonstrates that soil microbial legacies, shaped by decades of agricultural management and climate treatment, exert significant vultivar-specific effects on wheat drought responses and the wheat rhizosphere microbiome. By inoculating sterilized potting soil with soil microbes extracted from ambient- vs. future-climate plots under organic and conventional managed fields, we showed that the two cultivars, SU Fiete (drought-tolerant) and Nordkap (drought-sensitive), responded differently to microbial history. Across both cultivars, inoculated seeds germinated more rapidly and reached higher germination rates than non-inoculated controls, with conventional-ambient inocula accelerating Nordkap emergence. Nordkap tended to have higher DWC when inoculated with future-climate microbial extract under drought. SU Fiete, in contrast, experienced the highest DWC with ambient-climate inocula. Our findings further confirmed that the farming history of soil microbial extracts explained the largest proportion of change in the rhizosphere community structure for both bacteria and fungi. Across both cultivars, organic-derived inocula resulted in greater rhizosphere fungal Shannon diversity under drought and contributed the largest pool of unique rhizosphere taxa. Taken together, our results suggest that soil microbial legacies can either buffer or exacerbate plant drought responses, and that the direction and magnitude of these effects depend on the match between the plant breeding background and the soil microbial histories.

Microbes from conventional–ambient soils boost germination rate and time, whereas organic–future inocula delayed and suppressed seedling emergence. This result aligns with our previous study, where, based on agar-based evidence, we showed that conventional–ambient microbiomes enhanced seed emergence, highlighting that microbial legacy (farming, climate) modulates the seed establishment even in the absence of soil heterogeneity (Ornik *et al*. 2024). The direction of the microbial inoculation effect on seed germination is plausible given the breeding background of Nordkap (high-input, intensive yield environments) and SU Fiete (broad adaptability within fertilized, pesticide-amended systems). This breeding history aligns with our germination results that modern wheat varieties, bred to perform under conventional management, are “tuned” to benefit from microbial extracts molded by those very same practices, whereas organic-origin microbiomes, with their slower-cycling, confer less germination advantage. These results suggest that the compatibility between plant breeding strategies and microbial provenance acts as a key determinant of early seed establishment success. The effect of plant breeding on host microbiomes and genotype-dependent microbial assembly has been reported in multiple systems (Azarbad *et al*. 2018; Quiza *et al*. 2023). For instance, in perennial wheatgrass (*Thinopyrum intermedium*), even limited breeding cycles induced substantial shifts in seed endophyte composition, marked by reduced diversity and functional gene abundance (e.g., *nirK*, *nifH*). These microbiome changes, particularly the loss of ASVs related to *Actinobacteria*, *Alphaproteobacteria*, and *Bacilli*, were associated with genotypic selection and a narrowing of plant parental lines (Michl *et al*. 2024). Crop breeding efforts have begun to recognize the potential of microbiome-assisted selection, proposing the integration of microbial inheritance and stress responsiveness as traits under selection (Cernava 2024).

Drought adaptation is strongly linked to DWC, processes that determine how plants allocate resources between survival, growth, and reproduction (Cai *et al*. 2022). Water stress typically shifts allocation away from reproductive organs and toward vegetative tissues that sustain growth, with stalks alone able to redistribute up to 35% of their dry weight under stress (Cai *et al*. 2023). Drought-tolerant plants typically accumulate more structural biomass relative to water content, involving increased lignification, osmoprotectant accumulations, cuticle reinforcement, and root-to-shoot ratio adjustments (Bakker, Mommer and van Ruijven 2018; Zandalinas *et al*. 2018; Rivero *et al*. 2022). In our study, Nordkap maintained higher DWC when inoculated with future-climate microbial communities, suggesting a stress priming effect in which microbial legacies enhanced the efficiency of resource allocation under drought. By contrast, SU Fiete achieved higher DWC with ambient-climate microbes, indicating stabilized tissue biomass only when microbial inputs reflected less stressful climatic conditions. At GCEF, future-climate plots had ∼20% less summer rain, ∼10% more spring/autumn rain, and sustained warming (+0.55 °C daily; +1.1 °C night) (Schädler *et al*. 2019). Applied over years, this coupled moisture–temperature forcing acted as a directional filter on soil microbiomes, reshaping community composition (Wahdan *et al*. 2021; Ornik *et al*. 2024). Indeed, in Nordkap, genera such as *Steroidobacter, Chthoniobacter,* and *Achromobacter* showed a strong positive correlation with DWC under drought with future-climate inocula. These taxa are known to contribute to nitrogen cycling, carbon turnover, and stress signalling (Wang *et al*. 2018, 2023; Vázquez *et al*. 2024). Such (potential) microbial functional traits likely facilitated efficient resource partitioning and dry weight accumulation under drought.

Our results suggest that the same microbial history can confer an advantage or disadvantage depending on host genotype. This interpretation is consistent with the management × rhizosphere selection observed in long-term agricultural fields (maize-tomato rotations), where bacterial community composition differed by organic vs conventional management and by bulk vs rhizosphere, and the strength/direction of the rhizosphere effect (host filtering) depended on management history (Schmidt *et al*. 2019). Extending to climate legacies, an experiment with *Camelina sativa* showed that both cropping system and precipitation history defined the soil and root microbiome, with plant filtering subsequently acting on these legacies to shape the rhizosphere bacterial community (Barnes *et al*. 2025).

Agricultural practices have long been recognized for their role in shaping soil microbiomes (Hartmann *et al*. 2015; Azarbad *et al*. 2020; Azarbad 2022, 2024; Hartmann and Six 2022; Behr *et al*. 2024). In our study, organic-derived microbial inocula supported higher richness and Shannon diversity, particularly in Nordkap, and retained fungal diversity under drought stress, suggesting that low-input systems may harbor ecologically buffered microbiomes. Importantly, diversity effects were cultivar-dependent. SU Fiete, despite being more stable in control treatments, showed a reduction in bacterial richness under drought when paired with future-climate microbiomes, while Nordkap maintained bacterial diversity. The pattern that fungal diversity is buffered under organic legacies agrees with multi-site evidence that organic management increases fungal richness and network complexity and maintains AMF under stress (Wahdan *et al*. 2021; Nam, Lee and Choi 2023), while agricultural intensification simplifies networks and reduces keystone taxa (Banerjee *et al*. 2019) and soil functionality (van Rijssel *et al*. 2025).

Wheat rhizosphere contained a large ASVs core (>1,700 bacterial; >300 fungal ASVs). However, the rhizosphere of plants grown in soil inoculated with organic microbes contributed 442 bacterial and 70 fungal unique ASVs, compared with 381/48 under conventional and 73/60 in non-inoculated plants. These patterns are consistent with reports that organic systems harbour more heterogeneous soil microbiomes (Lupatini *et al*. 2017) and that management legacies shape plant-associated core microbiomes (Barnes *et al*. 2025). From an agricultural perspective, these results highlight the promise of organic systems as microbial reservoirs capable of buffering crops against climate stress. Indeed, Piton et al. (2021) showed that across European agroecosystems, conventional management increased the resistance of soil microbial biomass and extracellular enzyme activity to altered rain regimes, whereas ecological (organic-like) management enhanced post-stress recovery (Piton *et al*. 2021). However, another study revealed that both organic and conventional fields maintain distinct soil microbiomes under drought stress (i.e., identity is conserved), but neither system is inherently buffered against severe drought (Kost *et al*. 2024). In view of these contradictory results, multi-region experiments that replicate organic and conventional farming practices across contrasting climatic contexts, with practice intensity quantified on a common scale (e.g., tillage frequency and pesticide pressure), would be necessary to confirm our findings. In addition, further studies should consider applying drought–rewetting cycles to evaluate resistance vs. recovery, which is necessary to confirm resistance-resilience patterns.

Our results on the rhizosphere community structure confirmed the pattern we observed in diversity, where bacterial communities were shaped by complex multi-level interactions between farming, climate, drought, and cultivar, whereas fungi were governed only by farming. This microbial-specific divergence points to the fact that fungi carry the long-term imprint of agricultural management. Independent evidence from the same field platform reported that arbuscular mycorrhizal fungi (AMF) in wheat roots respond to both climate treatment and farming, and that organic management buffers AMF richness under future climate (Wahdan *et al*. 2021). It is also in line with long field trials in which bacterial communities were shaped by the interaction of management (organic vs conventional) and rhizosphere filtering, while fungal communities tracked management alone (Schmidt *et al*. 2019). More broadly, a large-scale field study revealed that intensive management and modern high-input cultivars divert carbon away from roots, thus reducing fine-root habitat for AMF and colonization under fertilization. As a consequence, agricultural intensification reduces AMF richness and biomass along with the loss of specialist fungi taxa (and a shift toward generalists) (Vahter *et al*. 2025). Another evidence is from farm-level surveys, which showed that organic farming legacy increases fungal richness and Shannon diversity and yields more complex community structures, whereas conventional legacies support simpler fungal and oomycete networks with lower phylogenetic diversity (Nam, Lee and Choi 2023). Collectively, these patterns may suggest that farming legacy shapes rhizosphere fungal diversity that persists across climate histories, while bacterial assembly carries additional filters, host genotype, and current drought exposure, that mediate short-timescale responses in the rhizosphere.

## Conclusion

Our study demonstrates that microbial legacy is an important complementary axis of crop responses to stress, alongside host genotype and abiotic stress. We showed that historical farming and climate conditions leave lasting soil microbial imprints that directly affect plant performance (germination, growth) and the plant rhizosphere. For agriculture, this means that breeding and management strategies must integrate microbial legacies explicitly, testing cultivar–microbiome compatibility under projected climate scenarios. Future work combining metagenomics and long-term field validation will be critical to identify the taxa and functions underpinning these legacies.

## Data availability

The data that support the findings of this study are available in the supplementary material of this article. Fastq files are deposited in the NCBI Sequence Read Archive (the BioProject accession PRJNA1085030).

## Acknowledgments

We would like to thank the research group at the Global Change Experimental Facility for providing us with the soil samples and access to the field. This work was funded by the Deutsche Forschungsgemeinschaft (DFG, German Research Foundation; project number: 523864000 to HA). Open Access funding provided by the Open Access Publication Fund of Philipps-Universität Marburg with support of the DFG.

## Author contributions

HA designed the study. MC planned and performed the greenhouse experiment. MS provided the soil and access to the GCEF field. BS performed the bioinformatic analyses, analyzed the data, and wrote the manuscript with the help of HA. All co-authors participated in reviewing and editing the final text. The authors declare no competing interests.

