## Supplementary material for "Soil microbial legacies and cultivar compatibility modulate the responses of wheat to drought": Tables

**Table 1.** ANOVA test results for the effects of Farming, Climate, Wheat Cultivar, Water Stress, and their interactions on fresh biomass and dry weight contents.

|  | **Fresh Biomass** | **Dry Weight Content** |
| --- | --- | --- |
| *R^2^* | 0.74 | 0.46 |
| *Farming* | *F=* 3.86 | *F=* 1.25 |
|  | *P=* 0.054 | *P=* 0.269 |
| *Climate* | *F=* 0.73 | *F=* 0.16 |
|  | *P=* 0.395 | *P=* 0.695 |
| *Cultivar* | *F=* 6.72 | *F=* 8.85 |
|  | *P=* **0.012** | *P=* **0.004** |
| *Water Stress* | *F=* 152.09 | *F=* 28.32 |
|  | *P***< 0.001** | *P***< 0.001** |
| *Farming × Climate* | *F=* 2.83 | *F=* 0.14 |
|  | *P=* 0.098 | *P=* 0.708 |
| *Farming × Cultivar* | *F=* 0.01 | *F=* 0.47 |
|  | *P=* 0.911 | *P=* 0.498 |
| *Climate × Cultivar* | *F=* 5.28 | *F=* 0.85 |
|  | *P=* **0.025** | *P=* 0.359 |
| *Farming × Water Stress* | *F=* 0.03 | *F=* 0.93 |
|  | *P=* 0.874 | *P=* 0.339 |
| *Climate × Water Stress* | *F=* 1.69 | *F=* 0.06 |
|  | *P=* 0.198 | *P=* 0.812 |
| *Cultivar × Water Stress* | *F=* 0.19 | *F=* 0.08 |
|  | *P=* 0.665 | *P=* 0.783 |
| *Farming × Climate × Cultivar* | *F=* 4.54 | *F=* 1.37 |
|  | *P=* **0.037** | *P=* 0.246 |
| *Farming × Climate × Water Stress* | *F=* 0.02 | *F=* 0.59 |
|  | *P=* 0.890 | *P=* 0.445 |
| *Farming × Cultivar × Water Stress* | *F=* 0.35 | *F=* 0.52 |
|  | *P=* 0.557 | *P=* 0.472 |
| *Climate × Cultivar × Water Stress* | *F=* 3.16 | *F=* 9.83 |
|  | *P=* 0.080 | *P=* **0.003** |
| *Farming × Climate × Cultivar × Water Stress* | *F=* 0.38 | *F=* 0.52 |
|  | *P=* 0.538 | *P=* 0.475 |

*ANOVA analysis excludes uninoculated controls of farming and climate.

Farming refers to the history of soil microbes extracted from conventional and organic agriculture.

Climate refers to history of soil microbes extracted from ambient and future climate under each farming.

Cultivars refers to Nordkap and SU Fiete

Bold values indicate statistical significance (P < 0.05).

**Table 2**. ANOVA test for the effects of farming and climate histories and their interactions on bacterial and fungal Shannon diversity and communities of rhizospheric soil extracts.

| **Factor** | **Bacteria** | | | | | | **Fungi** | | | | | | | |
| --- | --- | --- | --- | --- | --- | --- | --- | --- | --- | --- | --- | --- | --- | --- |
|  | **Richness** | | **Shannon diversity** | | **Community structure** | | **Richness** | | | **Shannon diversity** | | | **Community structure** | |
|  | **F**  **Value** | **P Value** | **F Value** | **P Value** | **F Value** | **P**  **Value** | **F**  **Value** | **P Value** | **F Value** | | **P Value** | **F Value** | | **P Value** |
| *Farming* | 3.375 | 0.071 | 0.415 | 0.522 | 3.780 | **0.001** | 1.812 | **0.001** | 3.902 | | 0.053 | 6.249 | | **0.001** |
| *Climate* | 0.397 | 0.531 | 2.689 | 0.106 | 3.720 | **0.002** | 0.003 | 0.958 | 0.025 | | 0.875 | 1.103 | | 0.312 |
| *Cultivar* | 1.039 | 0.312 | 0.191 | 0.664 | 1.149 | 0.251 | 1.518 | 0.223 | 0.678 | | 0.413 | 1.050 | | 0.346 |
| *Water Stress* | 14.110 | **0.000** | 0.410 | 0.524 | 2.462 | **0.003** | 0.182 | 0.671 | 3.378 | | 0.071 | 0.716 | | 0.626 |
| *Farming × Climate* | 4.961 | **0.029** | 0.918 | 0.342 | 1.774 | **0.022** | 0.047 | 0.830 | 0.230 | | 0.633 | 1.247 | | 0.254 |
| *Farming × Cultivar* | 2.437 | 0.123 | 9.179 | **0.004** | 2.415 | **0.005** | 1.610 | 0.209 | 0.150 | | 0.700 | 0.634 | | 0.680 |
| *Climate × Cultivar* | 0.160 | 0.691 | 0.242 | 0.625 | 1.257 | 0.174 | 0.708 | 0.403 | 0.103 | | 0.750 | 1.096 | | 0.331 |
| *Farming × Water Stress* | 0.000 | 0.995 | 0.602 | 0.441 | 1.104 | 0.273 | 0.339 | 0.563 | 1.913 | | 0.172 | 2.006 | | 0.086 |
| *Climate × Water Stress* | 0.021 | 0.886 | 1.181 | 0.281 | 0.991 | 0.460 | 0.634 | 0.429 | 1.680 | | 0.200 | 1.069 | | 0.337 |
| *Cultivar × Water Stress* | 2.845 | 0.097 | 0.005 | 0.945 | 0.801 | 0.732 | 0.385 | 0.537 | 0.403 | | 0.528 | 1.794 | | 0.112 |
| *Farming × Climate × Cultivar* | 0.442 | 0.508 | 0.229 | 0.634 | 1.360 | 0.122 | 0.035 | 0.851 | 1.124 | | 0.293 | 0.908 | | 0.427 |
| *Farming × Climate × Water Stress* | 0.343 | 0.560 | 0.127 | 0.723 | 0.938 | 0.506 | 0.019 | 0.890 | 2.275 | | 0.137 | 0.678 | | 0.655 |
| *Farming × Cultivar × Water Stress* | 0.012 | 0.913 | 0.640 | 0.427 | 1.246 | 0.170 | 1.773 | 0.188 | 1.509 | | 0.224 | 0.712 | | 0.597 |
| *Climate × Cultivar × Water Stress* | 8.017 | **0.006** | 1.088 | 0.301 | 1.628 | **0.036** | 2.725 | 0.104 | 0.464 | | 0.498 | 0.558 | | 0.752 |
| *Farming × Climate × Cultivar × Water Stress* | 2.713 | 0.104 | 1.853 | 0.178 | 2.253 | **0.009** | 0.894 | 0.348 | 0.006 | | 0.937 | 0.441 | | 0.845 |
| *ANOVA analysis excludes uninoculated controls of farming and climate.  Farming refers to the history of soil microbes extracted from conventional and organic agriculture.  Climate refers to history of soil microbes extracted from ambient and future climate under each farming.  Cultivars refers to Nordkap and SU Fiete  Bold values indicate statistical significance (P < 0.05). | | | | | | | | | | | | | | |
