## Supplementary information for "Soil microbial legacies and cultivar compatibility modulate the responses of wheat to drought"

**Table S1.** ANOVA test results for the effects of Farming, Climate, Cultivars, and their interactions on the germination rate.

|  | **Day 4** | **Day 5** |
| --- | --- | --- |
| *Farming* | *F=* 9.620 | *F=* 54.85 |
|  | ***P=* 0.002** | *P***< 0.001** |
| *Climate* | *F=* 11.541 | *F=* 18.00 |
|  | ***P=* 0.001** | *P***< 0.001** |
| *Cultivar* | *F=* 0.545 | *F=* 16.80 |
|  | *P=* 0.461 | *P***< 0.001** |
| *Farming × Climate* | *F=* 3.688 | *F=* 9.56 |
|  | *P=* 0.057 | *P=* **0.002** |
| *Farming × Cultivar* | *F=* 0.022 | *F=* 0.01 |
|  | *P=* 0.883 | *P=* 0.943 |
| *Climate × Cultivar* | *F=* 1.767 | *F=* 0.42 |
|  | *P=* 0.186 | *P=* 0.519 |
| *Farming × Climate × Cultivar* | *F=* 0.022 | *F=* 0.63 |
|  | *P=* 0.883 | *P=* 0.430 |

*ANOVA analysis excludes uninoculated controls of farming and climate.

Farming type include conventional and organic

Climate refers to ambient and future

Cultivars refers to Nordkap and SU Fiete

Bold values indicate statistical significance (P < 0.05).

**Table S2**. ANOVA test results for the effects of change in Farming, Climate, Wheat Cultivar, Water Stress, and their interactions on fresh biomass and dry weight contents.

|  | **Δ Fresh Biomass** | **Δ Dry Weight Content** |
| --- | --- | --- |
| *R^2^* | 0.74 | 0.46 |
| *Farming* | *F=* 2.28 | *F=* 1.00 |
|  | *P=* 0.14 | *P=* 0.32 |
| *Climate* | *F=* 0.43 | *F=* 0.13 |
|  | *P=* 0.51 | *P=* 0.72 |
| *Cultivar* | *F=* 1.92 | *F=* 0.12 |
|  | *P=* 0.17 | *P=* 0.73 |
| *Water Stress* | *F=* 0.07 | *F=* 0.41 |
|  | *P=* 0.79 | *P=* 0.52 |
| *Farming × Climate* | *F=* 0.01 | *F=* 0.11 |
|  | *P=* 0.93 | *P=* 0.74 |
| *Farming × Cultivar* | *F=* 0.01 | *F=* 0.38 |
|  | *P=* 0.93 | *P=* 0.54 |
| *Climate × Cultivar* | *F=* 3.12 | *F=* 0.69 |
|  | *P=* 0.08 | *P=* 0.41 |
| *Farming × Water Stress* | *F=* 0.01 | *F=* 0.75 |
|  | *P=* 0.32 | *P=* 0.39 |
| *Climate × Water Stress* | *F=*1.00 | *F=* 0.05 |
|  | *P=* 0.32 | *P=* 0.83 |
| *Cultivar × Water Stress* | *F=* 4.11 | *F=* 3.43 |
|  | *P=* **0.05** | *P=* 0.07 |
| *Farming × Climate × Cultivar* | *F=* 2.68 | *F=* 1.11 |
|  | *P=* 0.11 | *P=* 0.30 |
| *Farming × Climate × Water Stress* | *F=* 0.01 | *F=* 0.48 |
|  | *P=* 0.92 | *P=* 0.49 |
| *Farming × Cultivar × Water Stress* | *F=* 0.21 | *F=* 0.42 |
|  | *P=* 0.65 | *P=* 0.52 |
| *Climate × Cultivar × Water Stress* | *F=* 1.86 | *F=* 7.93 |
|  | *P=* 0.18 | *P=* **0.01** |
| *Farming × Climate × Cultivar × Water Stress* | *F=* 0.23 | *F=* 0.42 |
|  | *P=* 0.64 | *P=* 0.52 |

*ANOVA analysis excludes uninoculated controls of farming and climate.

Farming type include conventional and organic

Climate refers to ambient and future

Cultivars refers to Nordkap and SU Fiete

Bold values indicate statistical significance (P < 0.05).

**Table S3**. ANOVA test results for the effects of change in Farming, Climate, Wheat Cultivar, Water Stress, and their interactions on relative abundance of bacteria at the genus level. Only significant bacterial genera are shown.

| **Factors** | **Bacterial Genus** | | | | | | | | | | | | | | | |
| --- | --- | --- | --- | --- | --- | --- | --- | --- | --- | --- | --- | --- | --- | --- | --- | --- |
|  | ***Alkanibacter*** | ***Rhizobium*** | ***Bacillus*** | ***Burkholderia*** | ***Devosia*** | ***Gemmatimonas*** | ***Gryllotalpicola*** | ***Leifsonia*** | ***Mesorhizobium*** | ***Micropepsis*** | ***Nocardioides*** | **Other** | ***Paenibacillus*** | ***Phenylobacterium*** | ***Pseudolabrys*** | ***Sphingomonas*** |
| *Farming* | F = 0.981 | F = 1.045 | F = 0.737 | F = 0.171 | F = 1.332 | F = 0.123 | F = 2.25 | F = 12.261 | F = 4.084 | F = 2.679 | F = 0.014 | F = 0.976 | F = 8.265 | F = 9.116 | F = 6.256 | F = 1.477 |
|  | P = 0.326 | P = 0.311 | P = 0.394 | P = 0.680 | P = 0.253 | P = 0.727 | P = 0.139 | **P = 0.001** | **P = 0.048** | P = 0.107 | P = 0.905 | P = 0.327 | **P = 0.006** | **P = 0.004** | **P = 0.015** | P = 0.229 |
| *Climate* | F = 7.122 | F = 6.115 | F = 0.145 | F = 0.629 | F = 1.412 | F = 1.075 | F = 0.759 | F = 0.36 | F = 0.551 | F = 3.266 | F = 4.607 | F = 3.418 | F = 2.739 | F = 1.446 | F = 0.549 | F = 0.141 |
|  | **P = 0.009** | **P = 0.016** | P = 0.705 | P = 0.430 | P = 0.239 | P = 0.304 | P = 0.387 | P = 0.551 | P = 0.460 | P = 0.075 | **P = 0.036** | P = 0.069 | P = 0.103 | P = 0.234 | P = 0.462 | P = 0.708 |
| *Water stress* | F = 15.157 | F = 0.048 | F = 0.003 | F = 8.193 | F = 1.367 | F = 24.097 | F = 1.232 | F = 0.168 | F = 4.116 | F = 4.322 | F = 0.411 | F = 0.298 | F = 10.556 | F = 1.464 | F = 0.443 | F = 7.460 |
|  | **P = 0.000** | P = 0.828 | P = 0.959 | **P = 0.005** | P = 0.247 | **P = 0.000** | P = 0.271 | P = 0.683 | **P = 0.047** | **P = 0.042** | P = 0.524 | P = 0.587 | **P = 0.002** | P = 0.231 | P = 0.508 | **P = 0.008** |
| *Cultivar* | F = 1.200 | F = 0.119 | F = 2.118 | F = 5.732 | F = 3.004 | F = 0.307 | F = 0.934 | F = 1.272 | F = 1.304 | F = 0.187 | F = 1.352 | F = 0.010 | F = 1.608 | F = 0.559 | F = 0.001 | F = 0.302 |
|  | P = 0.277 | P = 0.731 | P = 0.150 | **P = 0.020** | P = 0.088 | P = 0.582 | P = 0.338 | P = 0.264 | P = 0.257 | P = 0.667 | P = 0.249 | P = 0.921 | P = 0.209 | P = 0.458 | P = 0.975 | P = 0.584 |
| *Farming × Climate* | F = 0.139 | F = 0.049 | F = 1.225 | F = 0.071 | F = 0.369 | F = 0.569 | F = 0.041 | F = 0.328 | F = 0.028 | F = 1.239 | F = 0.463 | F = 2.749 | F = 0.482 | F = 1.106 | F = 8.659 | F = 0.032 |
|  | P = 0.711 | P = 0.825 | P = 0.272 | P = 0.791 | P = 0.546 | P = 0.454 | P = 0.841 | P = 0.569 | P = 0.868 | P = 0.270 | P = 0.499 | P = 0.102 | P = 0.490 | P = 0.297 | **P = 0.005** | P = 0.859 |
| *Farming × Water stress* | F = 0.586 | F = 0.091 | F = 0.005 | F = 0.073 | F = 1.343 | F = 0.282 | F = 1.023 | F = 0.441 | F = 5.131 | F = 3.260 | F = 0.195 | F = 2.856 | F = 0.438 | F = 9.926 | F = 3.502 | F = 1.204 |
|  | P = 0.447 | P = 0.764 | P = 0.943 | P = 0.788 | P = 0.251 | P = 0.597 | P = 0.316 | P = 0.509 | **P = 0.027** | P = 0.076 | P = 0.660 | P = 0.096 | P = 0.510 | **P = 0.003** | P = 0.066 | P = 0.277 |
| *Climate × Water stress* | F = 0.032 | F = 0.198 | F = 0.713 | F = 2.346 | F = 1.878 | F = 0.482 | F = 0.953 | F = 0.539 | F = 4.490 | F = 0.017 | F = 0.026 | F = 0.279 | F = 0.075 | F = 0.982 | F = 5.088 | F = 0.323 |
|  | P = 0.859 | P = 0.658 | P = 0.402 | P = 0.131 | P = 0.175 | P = 0.490 | P = 0.333 | P = 0.466 | **P = 0.038** | P = 0.898 | P = 0.873 | P = 0.599 | P = 0.785 | P = 0.325 | **P = 0.028** | P = 0.572 |
| *Farming × Cultivar* | F = 2.411 | F = 0.701 | F = 1.282 | F = 2.596 | F = 3.597 | F = 0.155 | F = 0.034 | F = 0.062 | F = 0.024 | F = 0.087 | F = 7.784 | F = 9.496 | F = 1.547 | F = 2.583 | F = 1.083 | F = 0.665 |
|  | P = 0.125 | P = 0.406 | P = 0.262 | P = 0.112 | P = 0.062 | P = 0.695 | P = 0.855 | P = 0.804 | P = 0.878 | P = 0.769 | **P = 0.007** | **P = 0.003** | P = 0.218 | P = 0.113 | P = 0.301 | P = 0.418 |
| *Climate × Cultivar* | F = 22.564 | F = 0.257 | F = 0.048 | F = 0.00 | F = 1.237 | F = 0.524 | F = 4.259 | F = 1.106 | F = 0.068 | F = 1.548 | F = 0.215 | F = 0.144 | F = 0.007 | F = 1.294 | F = 2.637 | F = 0.154 |
|  | **P = 0.000** | P = 0.614 | P = 0.827 | P = 0.983 | P = 0.270 | P = 0.471 | **P = 0.043** | P = 0.297 | P = 0.795 | P = 0.218 | P = 0.644 | P = 0.706 | P = 0.935 | P = 0.260 | P = 0.109 | P = 0.697 |
| *Treatment × Cultivar* | F = 1.588 | F = 0.163 | F = 0.221 | F = 5.600 | F = 0.007 | F = 0.686 | F = 2.083 | F = 1.136 | F = 0.044 | F = 0.535 | F = 0.259 | F = 0.001 | F = 0.468 | F = 1.875 | F = 0.295 | F = 1.505 |
|  | P = 0.212 | P = 0.688 | P = 0.640 | **P = 0.021** | P = 0.933 | P = 0.410 | P = 0.154 | P = 0.290 | P = 0.835 | P = 0.467 | P = 0.6124 | P = 0.981 | P = 0.496 | P = 0.176 | P = 0.589 | P = 0.224 |
| *Farming × Climate × Water stress* | F = 0.249 | F = 0.899 | F = 2.023 | F = 0.764 | F = 0.083 | F = 4.903 | F = 0.632 | F = 2.071 | F = 4.544 | F = 1.819 | F = 0.084 | F = 0.008 | F = 0.180 | F = 3.063 | F = 1.741 | F = 0.184 |
|  | P = 0.620 | P = 0.347 | P = 0.160 | P = 0.385 | P = 0.775 | **P = 0.030** | P = 0.429 | P = 0.155 | **P = 0.037** | P = 0.182 | P = 0.772 | P = 0.929 | P = 0.673 | P = 0.085 | P = 0.192 | P = 0.669 |
| *Farming × Climate × Cultivar* | F = 0.085 | F = 1.042 | F = 0.000 | F = 0.023 | F = 0.093 | F = 0.100 | F = 2.001 | F = 0.117 | F = 0.698 | F = 0.073 | F = 6.745 | F = 3.230 | F = 0.092 | F = 3.522 | F = 0.182 | F = 0.417 |
|  | P = 0.771 | P = 0.311 | P = 0.985 | P = 0.880 | P = 0.762 | P = 0.752 | P = 0.162 | P = 0.734 | P = 0.407 | P = 0.788 | **P = 0.012** | **P = 0.077** | P = 0.763 | P = 0.065 | P = 0.672 | P = 0.521 |
| *Farming × Water stress × Cultivar* | F = 1.182 | F = 2.198 | F = 0.002 | F = 4.993 | F = 2.670 | F = 2.009 | F = 4.558 | F = 0.065 | F = 2.333 | F = 0.928 | F = 0.728 | F = 0.417 | F = 2.138 | F = 1.336 | F = 6.798 | F = 0.801 |
|  | P = 0.281 | P = 0.143 | P = 0.9614 | **P = 0.029** | P = 0.107 | P = 0.161 | **P = 0.037** | P = 0.799 | P = 0.132 | P = 0.339 | P = 0.397 | P = 0.521 | P = 0.149 | P = 0.252 | **P = 0.011** | P = 0.374 |
| *Climate × Water stress × Cultivar* | F = 3.426 | F = 0.071 | F = 0.410 | F = 0.040 | F = 0.222 | F = 0.299 | F = 0.288 | F = 1.161 | F = 0.62 | F = 0.243 | F = 4.130 | F = 6.912 | F = 0.097 | F = 1.753 | F = 4.435 | F = 0.260 |
|  | P = 0.069 | P = 0.7901 | P = 0.524 | P = 0.843 | P = 0.639 | P = 0.586 | P = 0.594 | P = 0.285 | P = 0.434 | P = 0.624 | **P = 0.046** | P = 0.011 | P = 0.757 | P = 0.190 | **P = 0.039** | P = 0.612 |
| *Farming × Climate × Water stress × Cultivar* | F = 4.482 | F = 2.683 | F = 5.162 | F = 0.047 | F = 18.900 | F = 2.224 | F = 1.435 | F = 3.086 | F = 2.241 | F = 5.946 | F = 3.533 | F = 2.047 | F = 0.166 | F = 6.218 | F = 5.406 | F = 0.460 |
|  | **P = 0.038** | P = 0.10 | **P = 0.027** | P = 0.829 | **P = 0.000** | P = 0.141 | P = 0.235 | P = 0.084 | P = 0.139 | **P = 0.018** | P = 0.065 | P = 0.157 | P = 0.685 | **P = 0.015** | **P = 0.023** | P = 0.501 |

**Table S4**. ANOVA test results for the effects of change in Farming, Climate, Wheat Cultivar, Water Stress, and their interactions on relative abundance of fungi at the genus level. Only significant fungal genera are shown.

| **Factors** | **Fungal Genus** | | | | | | | | | | | | | |
| --- | --- | --- | --- | --- | --- | --- | --- | --- | --- | --- | --- | --- | --- | --- |
|  | ***Alternaria*** | ***Blumeria*** | ***Cladosporium*** | ***Exophiala*** | **Fungi_gen_**  **Incertae_sedis** | ***Fusarium*** | ***Humicola*** | ***Pseudogymnoascus*** | ***Pyrenochaetopsis*** | ***Rhizopus*** | **Rozellomycota_gen_**  **Incertae_sedis** | ***Sarocladium*** | ***Talaromyces*** | ***Umbelopsis*** |
| *Farming* | F = 5.524 | F = 0.048 | F = 4.652 | F = 0.941 | F = 9.585 | F = 14.665 | F = 5.903 | F = 0.132 | F = 1.106 | F = 2.180 | F = 0.352 | F = 1.697 | F = 2.094 | F = 0.265 |
|  | **P = 0.022** | P = 0.828 | **P = 0.035** | P = 0.336 | **P = 0.003** | **P = 0.000** | **P = 0.018** | P = 0.717 | P = 0.297 | P = 0.145 | P = 0.555 | P = 0.196 | P = 0.153 | P = 0.609 |
| *Climate* | F = 0.099 | F = 0.457 | F = 0.582 | F = 0.187 | F = 0.633 | F = 7.021 | F = 0.510 | F = 0.642 | F = 0.000 | F = 1.210 | F = 0.378 | F = 0.386 | F = 0.025 | F = 2.110 |
|  | P = 0.754 | P = 0.502 | P = 0.448 | P = 0.667 | P = 0.429 | **P = 0.010** | P = 0.477 | P = 0.426 | P = 0.987 | P = 0.276 | P = 0.541 | P = 0.537 | P = 0.875 | P = 0.151 |
| *Water stress* | F = 0.153 | F = 6.787 | F = 0.003 | F = 3.463 | F = 0.242 | F = 1.009 | F = 0.693 | F = 0.137 | F = 0.867 | F = 4.353 | F = 5.596 | F = 0.019 | F = 4.092 | F = 3.138 |
|  | P = 0.697 | **P = 0.012** | P = 0.959 | P = 0.068 | P = 0.624 | P = 0.319 | P = 0.408 | P = 0.713 | P = 0.355 | **P = 0.041** | **P = 0.021** | P = 0.892 | **P = 0.047** | P = 0.081 |
| *Cultivar* | F = 0.103 | F = 0.189 | F = 2.048 | F = 3.265 | F = 0.002 | F = 0.011 | F = 0.566 | F = 1.102 | F = 3.462 | F = 0.154 | F = 2.494 | F = 6.373 | F = 0.118 | F = 2.749 |
|  | P = 0.749 | P = 0.665 | P = 0.157 | P = 0.076 | P = 0.966 | P = 0.916 | P = 0.454 | P = 0.298 | P = 0.070 | P = 0.696 | P = 0.119 | **P = 0.014** | P = 0.733 | P = 0.102 |
| *Farming × Climate* | F = 0.026 | F = 0.455 | F = 0.072 | F = 0.011 | F = 0.002 | F = 7.839 | F = 1.320 | F = 1.312 | F = 0.914 | F = 1.861 | F = 0.005 | F = 0.155 | F = 0.465 | F = 0.005 |
|  | P = 0.871 | P = 0.502 | P = 0.790 | P = 0.915 | P = 0.965 | **P = 0.007** | P = 0.255 | P = 0.257 | P = 0.343 | P = 0.177 | P = 0.941 | P = 0.695 | P = 0.498 | P = 0.945 |
| *Farming × Water stress* | F = 1.204 | F = 0.548 | F = 1.269 | F = 1.865 | F = 0.094 | F = 0.067 | F = 0.131 | F = 0.783 | F = 0.005 | F = 4.893 | F = 0.456 | F = 1.391 | F = 2.217 | F = 0.020 |
|  | P = 0.277 | P = 0.462 | P = 0.264 | P = 0.177 | P = 0.760 | P = 0.797 | P = 0.719 | P = 0.380 | P = 0.947 | **P = 0.031** | P = 0.502 | P = 0.243 | P = 0.142 | P = 0.889 |
| *Climate × Water stress* | F = 1.043 | F = 0.000 | F = 0.024 | F = 0.291 | F = 2.619 | F = 1.108 | F = 5.295 | F = 0.324 | F = 1.284 | F = 2.495 | F = 0.326 | F = 3.151 | F = 0.002 | F = 0.764 |
|  | P = 0.311 | P = 0.992 | P = 0.878 | P = 0.591 | P = 0.111 | P = 0.297 | **P = 0.025** | P = 0.571 | P = 0.262 | P = 0.120 | P = 0.570 | P = 0.081 | P = 0.968 | P = 0.385 |
| *Farming × Cultivar* | F = 0.894 | F = 1.514 | F = 0.275 | F = 0.002 | F = 0.047 | F = 0.054 | F = 1.658 | F = 0.093 | F = 0.744 | F = 0.916 | F = 0.307 | F = 1.647 | F = 0.136 | F = 0.004 |
|  | P = 0.348 | P = 0.223 | P = 0.602 | P = 0.961 | P = 0.829 | P = 0.818 | P = 0.203 | P = 0.762 | P = 0.391 | P = 0.342 | P = 0.582 | P = 0.204 | P = 0.714 | P = 0.949 |
| *Climate × Cultivar* | F = 0.087 | F = 2.474 | F = 3.613 | F = 0.017 | F = 1.382 | F = 0.000 | F = 0.813 | F = 2.232 | F = 0.001 | F = 0.522 | F = 0.017 | F = 0.410 | F = 1.819 | F = 4.242 |
|  | P = 0.769 | P = 0.121 | P = 0.062 | P = 0.898 | P = 0.244 | P = 0.985 | P = 0.371 | P = 0.140 | P = 0.993 | P = 0.473 | P = 0.897 | P = 0.524 | P = 0.182 | **P = 0.044** |
| *Treatment × Cultivar* | F = 0.501 | F = 1.769 | F = 1.055 | F = 8.159 | F = 0.737 | F = 0.018 | F = 0.000 | F = 0.231 | F = 0.772 | F = 0.024 | F = 3.370 | F = 0.014 | F = 0.119 | F = 1.342 |
|  | P = 0.482 | P = 0.188 | P = 0.308 | **P = 0.006** | P = 0.394 | P = 0.895 | P = 0.984 | P = 0.633 | P = 0.383 | P = 0.878 | P = 0.071 | P = 0.908 | P = 0.731 | P = 0.251 |
| *Farming × Climate × Water stress* | F = 0.003 | F = 0.332 | F = 0.840 | F = 0.486 | F = 0.634 | F = 0.757 | F = 4.332 | F = 4.343 | F = 4.217 | F = 0.756 | F = 0.358 | F = 4.333 | F = 0.156 | F = 0.391 |
|  | P = 0.900 | P = 0.567 | P = 0.363 | P = 0.488 | P = 0.429 | P = 0.388 | **P = 0.041** | **P = 0.041** | **P = 0.044** | P = 0.388 | P = 0.552 | **P = 0.041** | P = 0.694 | P = 0.534 |
| *Farming × Climate × Cultivar* | F = 0.003 | F = 0.954 | F = 0.390 | F = 0.099 | F = 0.017 | F = 0.432 | F = 0.048 | F = 0.016 | F = 0.818 | F = 0.591 | F = 1.075 | F = 0.161 | F = 1.485 | F = 0.081 |
|  | P = 0.958 | P = 0.339 | P = 0.535 | P = 0.754 | P = 0.896 | P = 0.513 | P = 0.828 | P = 0.901 | P = 0.369 | P = 0.445 | P = 0.304 | P = 0.690 | P = 0.228 | P = 0.776 |
| *Farming × Water stress × Cultivar* | F = 0.209 | F = 0.532 | F = 0.157 | F = 2.160 | F = 1.964 | F = 0.049 | F = 0.220 | F = 0.895 | F = 0.005 | F = 0.534 | F = 0.189 | F = 1.415 | F = 0.186 | F = 0.284 |
|  | P = 0.649 | P = 0.469 | P = 0.693 | P = 0.147 | P = 0.166 | P = 0.825 | P = 0.641 | P = 0.34 | P = 0.945 | P = 0.468 | P = 0.666 | P = 0.239 | P = 0.668 | P = 0.596 |
| *Climate × Water stress × Cultivar* | F = 0.058 | F = 0.063 | F = 0.247 | F = 1.436 | F = 2.157 | F = 0.069 | F = 1.800 | F = 0.331 | F = 0.904 | F = 3.413 | F = 0.034 | F = 3.117 | F = 1.071 | F = 2.836 |
|  | P = 0.811 | P = 0.803 | P = 0.621 | P = 0.235 | P = 0.147 | P = 0.794 | P = 0.185 | P = 0.567 | P = 0.345 | P = 0.070 | P = 0.854 | P = 0.082 | P = 0.305 | P = 0.097 |
| *Farming × Climate × Water stress × Cultivar* | F = 2.97 | F = 1.330 | F = 0.004 | F = 0.036 | F = 0.046 | F = 0.066 | F = 0.737 | F = 1.585 | F = 3.595 | F = 0.092 | F = 0.898 | F = 4.079 | F = 1.103 | F = 0.048 |
|  | P = 0.090 | P = 0.253 | P = 0.952 | P = 0.849 | P = 0.831 | P = 0.797 | P = 0.394 | P = 0.213 | P = 0.063 | P = 0.762 | P = 0.347 | **P = 0.050** | P = 0.300 | P = 0.827 |

**Table S5.** Spearman correlation between DWC and abundancies of significant fungal genera in the rhizosphere of wheat under ambient and future climate conditions

| **Genus** | **Ambient** | **Future** |
| --- | --- | --- |
| **Nordkap** | | |
| *Cladosporium* | -0.664^*^ | 0.1732 |
| **SU Fiete** | | |
| *Talaromyces* | -0.383 | 0.731^*^ |
| *Fusarium* | -0.035 | -0.731^*^ |
| *Oidiodendron* | 0.643^*^ | 0.050 |
| Fungi_gen_Incertae_sedis | 0.731^*^ | -0.453 |
| *Pseudogymnoascus* | 0.811^**^ | 0.108 |
| *Cladosporium* | 0.594 | -0.661^*^ |
| *Alternaria* | 0.140 | -0.661^*^ |
| *Gymnostellatospora* | 0 | -0.643^*^ |

*: p < 0.05

**: p < 0.01

***: p < 0.001

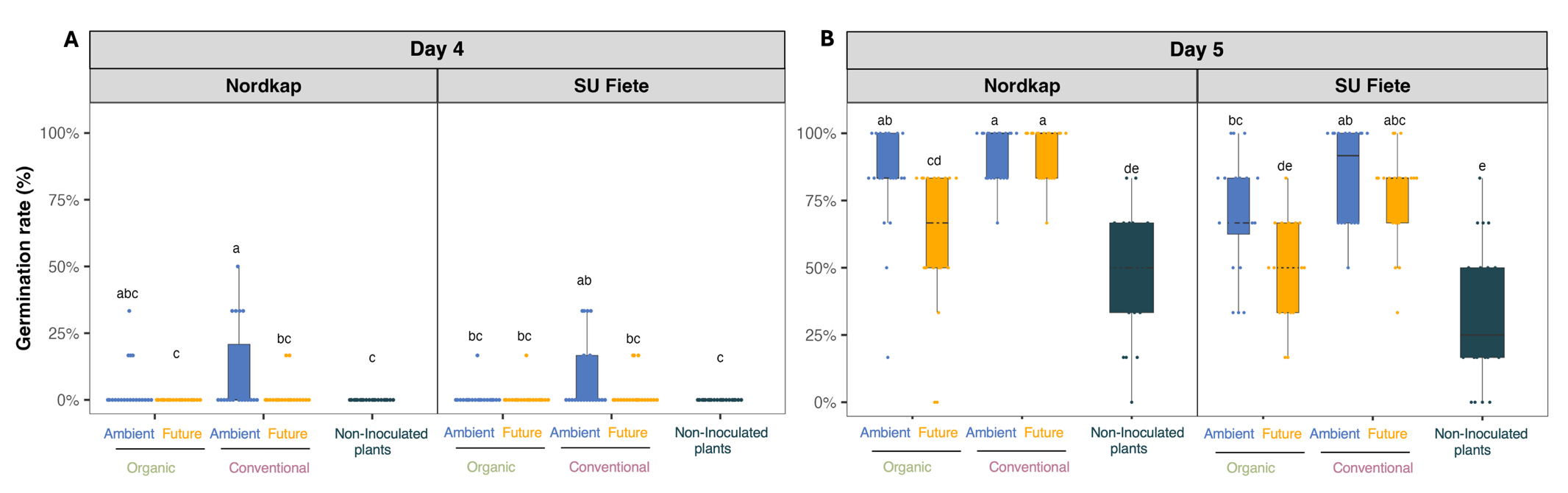

**Fig S1.** Germination rates and germination times of wheat seeds inoculated with soil microbial extract from different farming (organic vs. conventional) and climate (future vs. ambient) histories and non-inoculated control plants. Letters represent significant differences based on post hoc Tukey’s test.

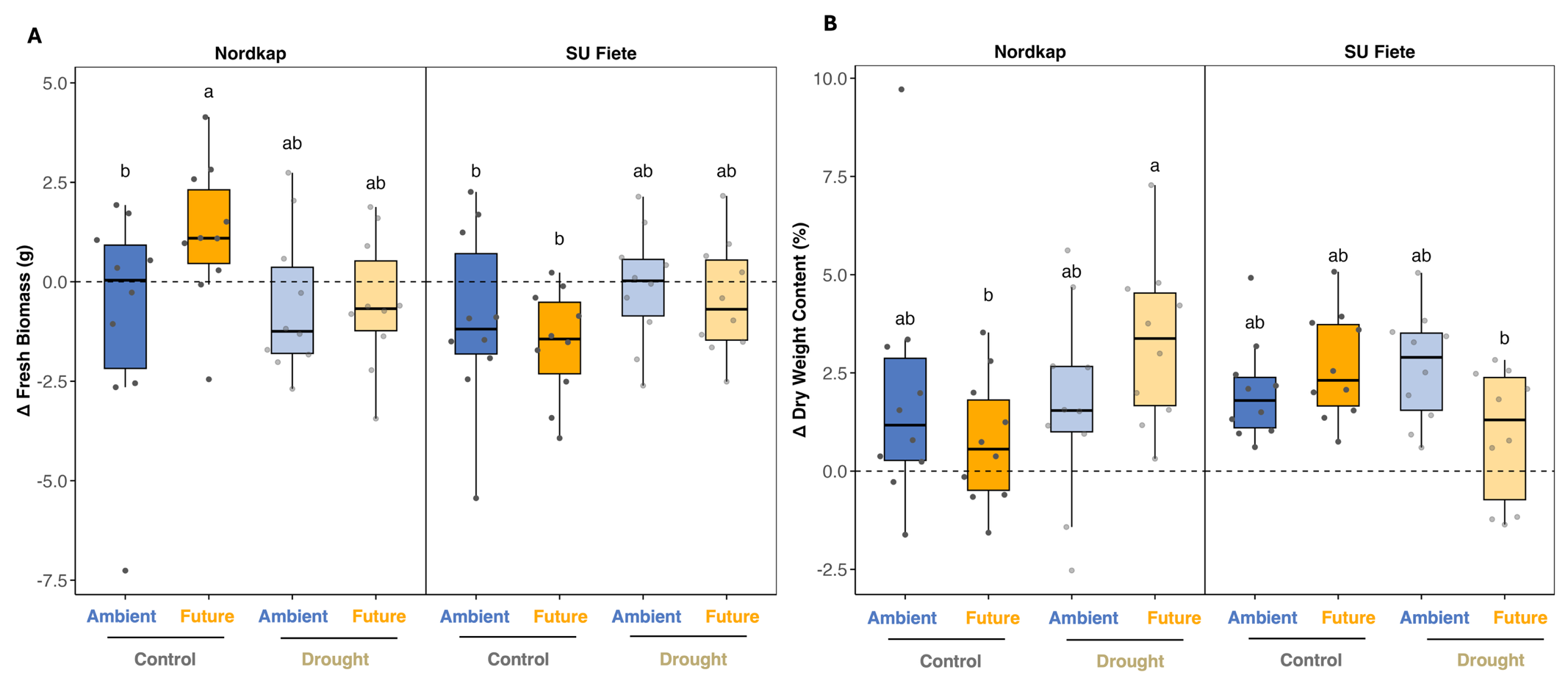

**Fig S2.** Change in aboveground biomass of wheat plants inoculated with soil microbial extract: (A) fresh biomass, (B) dry weight content. The dotted line indicates the corresponding biomass of non-inoculated plants. The dashed horizontal line represents no difference from the non‐inoculated control. A positive Δ indicates that inoculated plants produced more fresh biomass and DWC than their non‐inoculated counterparts, whereas a negative Δ represents a reduction. Letters represent significant differences.
